# Continental-scale survey of *Treponema pallidum* in African nonhuman primates: expanding the range to the Mediterranean basin

**DOI:** 10.64898/2026.09.02.748841

**Authors:** Clément Labarrere, Charles Kumakamba, Aboubacar Diakité, Aku Essenou Elomou, Mbayang Faye, Georges Diatta, Laia Dotras, Manuel Llana, Jean Akiana, Jean-Jacques Muyembe-Tamfum, R. Adriana Hernandez-Aguilar, Abdelbarie Baroudi, Younes Laidoudi, Bernard Davoust, Cheikh Sokhna, Florence Fenollar, Inestin Amona, Oleg Mediannikov

## Abstract

**Background:** *Treponema pallidum*, the causative agent of yaws, bejel, and syphilis, is a significant pathogen affecting both humans and nonhuman primates (NHPs). While its presence is documented in several African NHP species, the continental-scale distribution, ecological drivers, and status in North African populations remain poorly understood. We conducted a large-scale, non-invasive survey to investigate the circulation of *T. pallidum* across African NHP populations and evaluate geographical and ecological patterns of exposure.

**Results:** A total of 1,696 fecal samples from 11 wild NHP species collected across five African countries were screened for anti-*T. pallidum* antibodies using an automated chemiluminescent Simple Western assay, and fecal and dipteran samples were analyzed by molecular methods. Widespread seropositivity was observed across all African sites and multiple primate taxa, including great apes and cercopithecids. We provide the first evidence of *T. pallidum* antibodies in Barbary macaques (*Macaca sylvanus*) from North Africa, extending the known distribution of the pathogen to the Mediterranean basin. In West Africa, chimpanzees (*Pan troglodytes verus*) from Senegal and Guinea exhibited particularly high seroprevalence, confirming this region as a major endemic hotspot. In Central Africa, substantial circulation was documented in *Cercopithecus wolfi*, *Cercopithecus ascanius*, and western lowland gorillas (*Gorilla gorilla gorilla*). Arid/savanna biomes showed significantly higher exposure than Mediterranean and tropical rainforest biomes. No *T. pallidum* DNA was detected in fecal or dipteran samples.

**Conclusion:** Our findings demonstrate widespread circulation of *T. pallidum* among African NHPs, reveal its presence in North African primate populations for the first time. The observed high seroprevalence, coupled with significant variations across species and geographical regions, suggests the existence of multiple transmission pathways adapted to specific host behaviors and environmental constraints. These results support the existence of a broad multihost sylvatic reservoir and provide new insights into the ecology of *T. pallidum*, with important implications for wildlife health, One Health surveillance, and yaws eradication efforts.

## INTRODUCTION

Treponematoses, caused by the bacterium *Treponema pallidum*, represent a significant global health burden (1,2). While historically categorized into two distinct entities –non-venereal forms (yaws and bejel) and the venereal form (syphilis)– recent genomic and epidemiological evidence has considerably expanded our understanding of the ecology and host range of *T. pallidum*. Regarding *T. pallidum* subsp. *pertenue* (TPE) in particular, a major paradigm shift is currently underway, driven by the increasing documentation of this infection in nonhuman primates (NHPs) (3–7). These discoveries challenge the long-held view of yaws as a strictly human disease and highlights its broad host range across sub-Saharan Africa.

These infections, which mirror the clinical manifestations of human yaws and syphilis (including ulcerative skin lesions, bone deformities, and severe orofacial or genital ulcerations that can lead to lethal outcomes) raise critical questions regarding wildlife health, conservation, and the potential for zoonotic spillover (8,9).

To date, simian treponematosis has been reported in numerous NHP populations across sub-Saharan Africa, with seroprevalence reaching 70–80% in some communities (4,6,10). However, these data are almost exclusively derived from regions currently or historically endemic for yaws. This narrow focus has created a significant geographic and ecological gap in our understanding. Consequently, the status of NHPs in arid or Mediterranean climates, such as those of North Africa where bejel was historically endemic, remains entirely unexplored (2). Determining whether these host populations are restricted to tropical belts or extend to other geographical zones is essential for a comprehensive mapping of *T. pallidum* ecology and may provide critical insights into the evolutionary history of non-venereal treponematoses.

Despite the high prevalence of treponematoses in NHPs and the demonstrated human pathogenicity of the only isolated simian strain to date, namely the Fribourg-Blanc strain – which is nearly genetically identical to human yaws strains– no direct cross-species transmission has been formally documented in a natural setting (11,12). However, the persistence of these pathogens in diverse animal host populations poses a latent threat to global eradication efforts, thereby necessitating a One Health approach that integrates human, animal, and environmental surveillance (13).

Beyond direct contact, the epidemiological framework of treponematoses is further complicated by the potential role of environmental vectors. Recent evidence suggests that hematophagous or necrophagous dipterans may contribute to the environmental persistence and potential mechanical transmission of *T. pallidum*, particularly in areas of close contact between NHPs. Recent molecular studies have detected *T. pallidum* DNA in 17–24% of wild-caught flies in Tanzania, with sequences clustering alongside *T. pallidum* subsp. *pallidum* (TPA) and *T. pallidum* subsp. *endemicum* (TEN) strains found in humans, as well as TPE strains infecting both humans and NHPs (14). Similarly, in Côte d’Ivoire, 6.1% of flies collected in proximity to infected sooty mangabey (*Cercocebus atys atys*) groups tested positive for *T. pallidum* (15). These findings suggest that dipterans may act as environmental conduits for the mechanical dissemination of *T. pallidum*, highlighting the necessity of integrating ecological and entomological factors into a comprehensive understanding of *T. pallidum* transmission dynamics.

In this context, we conducted a large-scale investigation to assess the prevalence of *T. pallidum* infection in NHPs across Africa. To this end, fecal samples were collected non-invasively, and fecal serology was performed to determine the serological status of the primates. Our study sites include both yaws-endemic regions in sub-Saharan Africa and previously uncharacterized Mediterranean habitats in North Africa. In parallel, we evaluated the presence of *T. pallidum* DNA in dipterans collected in proximity to primate populations. By including both endemic and previously unexplored region, this study provides updated insights into the epidemiology of treponemal infections and contributes to a more comprehensive understanding of their transmission dynamics.

## MATERIALS AND METHODS

### Sample collection

NHP fecal samples were collected across five African countries (Figure 1). Sampling was conducted in Algeria (2018 and 2025), the Democratic Republic of the Congo (DRC; 2023), the Republic of Guinea (2024), the Republic of the Congo (2019–2025), and Senegal (2021–2025). In Algeria, samples were collected from several localities in the northern part of the country, including Chréa National Park, Béjaïa, Tikjda, Médéa, and Boumerdès. In the DRC, sampling was performed in the Mabali Forest Reserve (Équateur Province). In the Republic of Guinea, samples were collected from three protected areas: Haut Niger National Park, Moyen-Bafing National Park, and the Diecké Forest Reserve. In the Republic of the Congo, samples were obtained from wild and semi-captive primates across four sites: Odzala-Kokoua National Park, Lesio-Louna Gorilla Nature Reserve, Tchimpounga Chimpanzee Sanctuary, and Kouilou-Konkouati National Park. In Senegal, sampling was conducted in Dindefelo Community Nature Reserve, Djoudj National Park, and Mako National Park.

**Figure 1.**
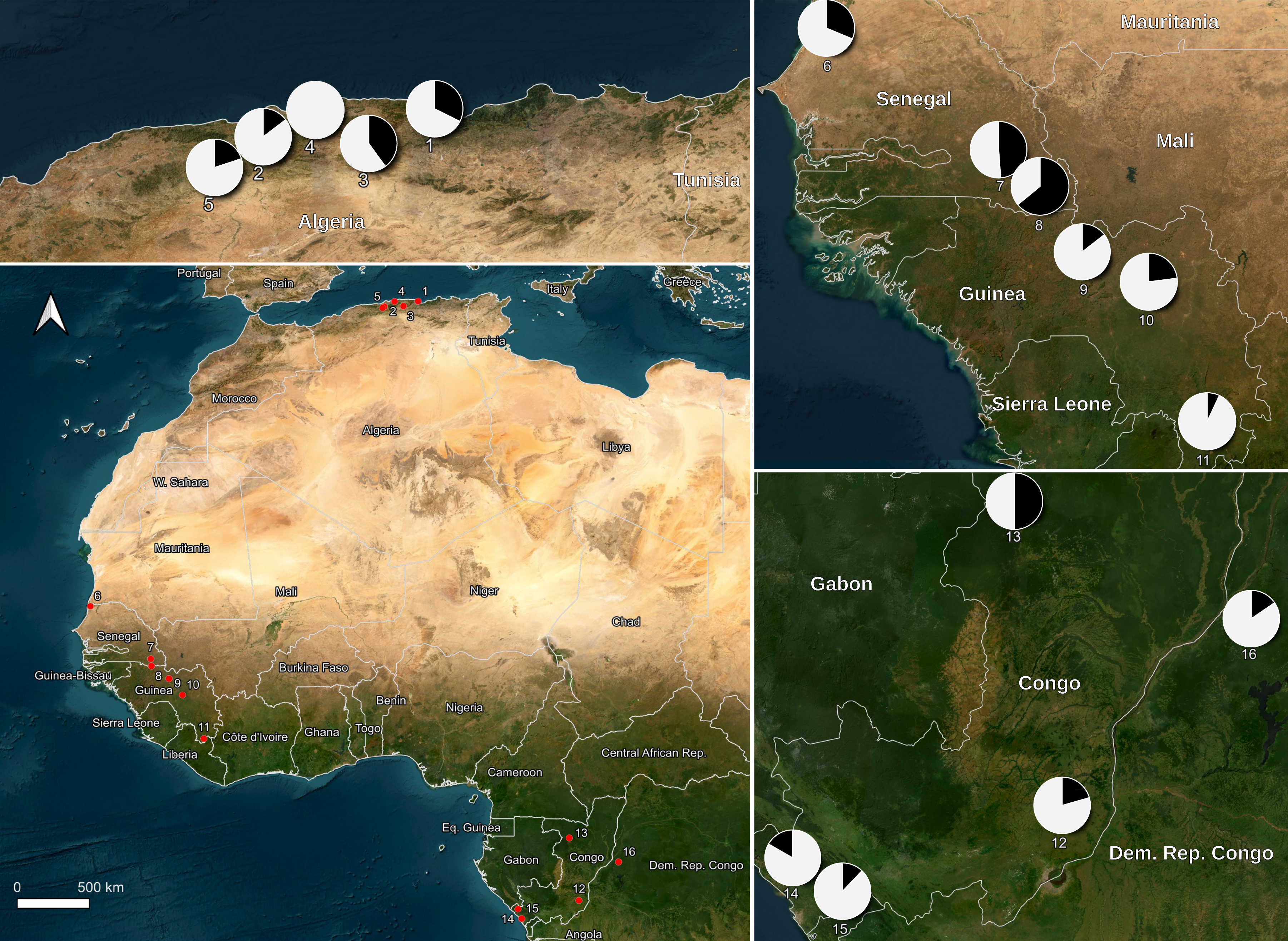
Geographic distribution and seroprevalence of *Treponema pallidum* in African nonhuman primates. The main map shows the 13 taxa from 12 sampled species across 16 study sites. Insets provide detailed views for Algeria (top left), Senegal and Guinea (right), and the Congo Basin (bottom right). Pie charts represent the proportion of seropositive (black) and seronegative (white) individuals; numbers correspond to site IDs: Algeria: 1 (Béjaïa, seroprevalence 32.4%), 2 (Chréa NP, 14.7%), 3 (Tikjda NP, 40%), 4 (Boumerdès, 0%), 5 (Médéa, 20%); Senegal: 6 (Djoudj, 31.3%), 7 (Mako, 48.8%), 8 (Dindefelo CNR, 64%); Guinea: 9 (Moyen-Bafing NR, 14.5%), 10 (Haut-Niger NP, 23.0%), 11 (Diecké FR, 7.0%); the Republic of the Congo: 12 (Lesio-Louna NR, 20.6%), 13 (Odzala-Kokoua NP, 50%), 14 (Konkouati-Douli NP, 83.3%), 15 (Tchimpounga Sanctuary, 12.1%); DRC: 16 (Équateur Province, 15.5%). Maps were created using QGIS with ESRI Satellite and World Boundaries basemaps. (NP: National Park; CNR: Community Nature Reserve, FR: Forest Reserve; NR: Natural Reserve; DRC: the Democratic Republic of the Congo).

Samples collected in Dindefelo were obtained within the framework of a long-term primate monitoring program conducted by the Jane Goodall Institute.

Control samples were collected from NHPs housed in two French zoological parks: Beauval and La Vallée des Singes. All fecal samples were stored in sterile containers at ≤ 4 °C during transport and frozen at −20 °C until analysis. Additionally, dipteran insects were collected in Senegal (2025) using sweep nets and biconical traps near nesting sites.

### Molecular methods

DNA was extracted from approximatively 20 mg of fecal material using the KingFisher Flex system (Thermo Fisher Scientific, Illkirch-Graffenstaden, France) combined with the NucleoMag Pathogen Kit (Macherey-Nagel, Hoerdt, France) following a 4-hour proteinase K incubation at 56 °C. Extracted DNA was stored at −20 °C until further analysis.

Host species identification was performed by amplification and sequencing of the mitochondrial 16S rRNA gene using the universal primers 16Sbr (5′-CCGGTCTGAACTCAGATCACGT-3′) and 16Sar (5′-CGCCTGTTTATCAAAAACAT-3′) (16). PCR reactions were carried out on a MiniAmp Thermal Cycler (Thermo Fisher Scientific). The reaction mixture consisted of 5 µL of template DNA, 12.5 µL of AmpliTaq Gold 360 Master Mix (Thermo Fisher Scientific), 0.75 µL of each primer (20 µM), and 6 µL of DNA/RNA-free ultrapure distilled water, for a final volume of 25 µL. The amplification program included an initial denaturation step at 95 °C for 15 min, followed by 35 cycles of denaturation at 95 °C for 30 s, annealing at 58 °C for 30 s, and extension at 72 °C for 1 min, with a final extension at 72 °C for 7 min. Following purification using Macherey-Nagel plates (EURL, Hoerdt, France), amplicons were sequenced using the BigDye Terminator Cycle Sequencing Kit on a 3500 Genetic Analyzer (Applied Biosystems).

All fecal samples were screened for *T. pallidum* DNA using a real-time PCR assay targeting the *polA* gene (16). Real-time PCR assays were performed on a CFX Opus Real-Time PCR System (Bio-Rad Laboratories, Hercules, USA). Each reaction contained 10 µL of LightCycler 480 Probes Master (Roche, Basel, Switzerland), 3.5 µL of DNase/RNase-free water, 0.5 µL of primers (20 µM), 0.5 µL of probe (5 µM), and 5 µL of extracted DNA, for a final reaction volume of 20 µL. The amplification protocol consisted of an initial denaturation at 95 °C for 10 min, followed by 45 cycles of 95 °C for 10 s and 60 °C for 30 s.

DNA extraction from dipteran insects and their transport medium was performed as previously described (14). Insects were identified by *COI* sequencing and screened for *T. pallidum* via *polA* qPCR, following the protocol used for fecal samples (18).

### Serological procedure

#### Protein extraction

For each sample, approximatively 1 g of fecal material were homogenized in 5 mL of extraction buffer composed of 1X DPBS (Dulbecco’s Phosphate-Buffered Saline; Thermo Fisher Scientific) supplemented with EDTA-free protease inhibitor tablets (cOmplete Mini, Merck; Reference: 11836170001; one tablet dissolved in 50 mL of DPBS). The feces buffer mixture was vortexed for 1 min and then incubated for 60 min at room temperature on a roller mixer at 8 rpm. Samples were subsequently centrifuged at 2,000 x g for 15 min at 4 °C. The supernatant was collected and filtered using a syringe-mounted 0.22 µm filter. After filtration, the protein fraction was lyophilized for 24 h at −80 °C and resuspended in 200 µL of ultrapure distilled water (Thermo Fisher Scientific). The lyophilizate was allowed to resuspend for 15 min at 4 °C. The entire resuspended volume was then subjected to ultrafiltration using Amicon® Ultra-4 centrifugal filter units with a molecular weight cut-off of 50 kDa (Reference: UFC805096; Sigma Aldrich Chimie, Saint-Quentin-Fallavier, France) following the manufacturer’s protocol. Protein extracts were stored at −20 °C until use.

#### Automated Western blot

Anti-*T. pallidum* antibodies were detected using the Jess^TM^ Simple Western platform (ProteinSimple, San Jose, USA), an automated system that couples capillary-based size separation with an integrated solid-phase nano-immunoassay. A recombinant antigen (70 kDa, Reference: ABIN934736, Antibodies-Online GmbH, Aachen, Germany) derived from the fusion of three major *T. pallidum* proteins (TpN15, TpN17, TpN47) commonly used in the serological diagnosis of treponematoses was employed (19). Western blot analyses were conducted using the 12–230 kDa separation module (SM-W004, ProteinSimple), following the manufacturer’s instructions. The recombinant antigen was loaded at a concentration of 0.2 µg/µL. Antigen–antibody complexes were detected using goat anti-human IgA, IgG, and IgM secondary antibodies conjugated to horseradish peroxidase (1:500 dilution; Jackson ImmunoResearch, Ely, UK). Chemiluminescent detection was carried out using the peroxide/luminol-S substrates provided by the manufacturer (ProteinSimple).

Digital images were acquired and analyzed based on chemiluminescence peak intensity and signal-to-noise (S/N) ratio using Compass for SW software (version 4.1.0, Protein Simple).

### Data analysis

Nucleotide sequences were assembled using ChromasPro (v1.7) and compared with reference sequences available in GenBank using the BLASTn algorithm (https://blast.ncbi.nlm.nih.gov/Blast.cgi).

For the Jess^TM^ automated Western blot, seropositivity was determined based on the S/N ratio at the specific molecular weight of the recombinant antigen (70 kDa). To establish a statistically robust threshold, fecal samples from NHPs housed in two French zoological parks were used as negative controls. These animals were under continuous veterinary surveillance and confirmed negative for treponemal infection. The cut-off for seropositivity was defined as the mean S/N ratio of these negative controls plus three standard deviations (mean + 3 SD).

This conservative threshold was selected to maximize diagnostic specificity and mitigate the risk of false-positive results inherent to the complexity of the fecal matrix. Samples exhibiting an S/N ratio strictly greater than this value were classified as seropositive.

Statistical analyses were performed using R version 4.2.1 (R Foundation for Statistical Computing, Vienna, Austria). To evaluate the impact of environmental and host-related factors on treponemal circulation, sampling sites were categorized into three distinct biomes (arid/savanna, tropical rainforest, and Mediterranean). Differences in seroprevalence between biomes and host taxa (great apes vs. cercopithecids) were assessed using *n* x 2 contingency tables and the Pearson’s Chi-square ( ^2^) test. For pairwise comparisons between specific sites or species, a two-tailed Fisher’s exact test was employed. The strength of the association between host type and seropositivity was quantified using Odds Ratios (OR) with their corresponding 95% confidence intervals (95% CI).

Temporal stability of seroprevalence for chimpanzees at the longitudinal site (Dindefelo) was tested using a χ^2^ test for trend. All proportions were tested at a critical probability level of 0.05, and a *p*-value < 0.05 was considered statistically significant.

## RESULTS

### Sample collection and host identification

In Africa, a total of 1,696 fecal samples were collected across five countries, representing a diverse range of biomes (Figure 1). In France, 30 samples were obtained from ZooParc de Beauval and four from La Vallée des Singes, when the primates underwent routine veterinary health examinations and served as seronegative controls in this study and served as seronegative controls in this study (10). In Africa, sampling spanned diverse ecosystems, from the dry savannas of Senegal to the tropical rainforests of the Republic of the Congo and the Mediterranean habitats of North Africa. In Algeria, 74 samples were collected in 2018 and 40 in 2025. In the DRC, 569 samples were collected in 2023. In the Republic of Guinea, 500 samples were collected in 2024. In the Republic of the Congo, 24 samples were collected in 2019, 97 in 2021, and 20 in 2025. In Senegal, 36 samples were collected in 2021, 124 in 2024, and 178 in 2025.

Host species identification via mitochondrial 16S rRNA sequencing confirmed the presence of 13 different NHP taxa belonging to 12 species (Table 1). In the two zoological parks in France, all fecal samples (n=34) were collected from *Pan troglodytes verus*. In Algeria, all samples (n=114) were confirmed to originate from Barbary macaques (*Macaca sylvanus*). In the DRC, all samples were collected from non-hominoid primates, predominantly *Cercopithecus ascanius* (n=354), *Lophocebus aterrimus* (n=148), and *C. wolfi* (n=61) (20). In the Republic of Guinea, all fecal samples (n=500) were collected from *P. t. verus*. In the Republic of the Congo, 50 samples originated from *Gorilla gorilla gorilla* and 91 from *P. t. troglodytes*. In Senegal, 142 samples were obtained from *P. t. verus*, 124 from *Papio papio*, 67 from *Erythrocebus patas* and 5 from *Chlorocebus sabaeus*.

**Table 1.**
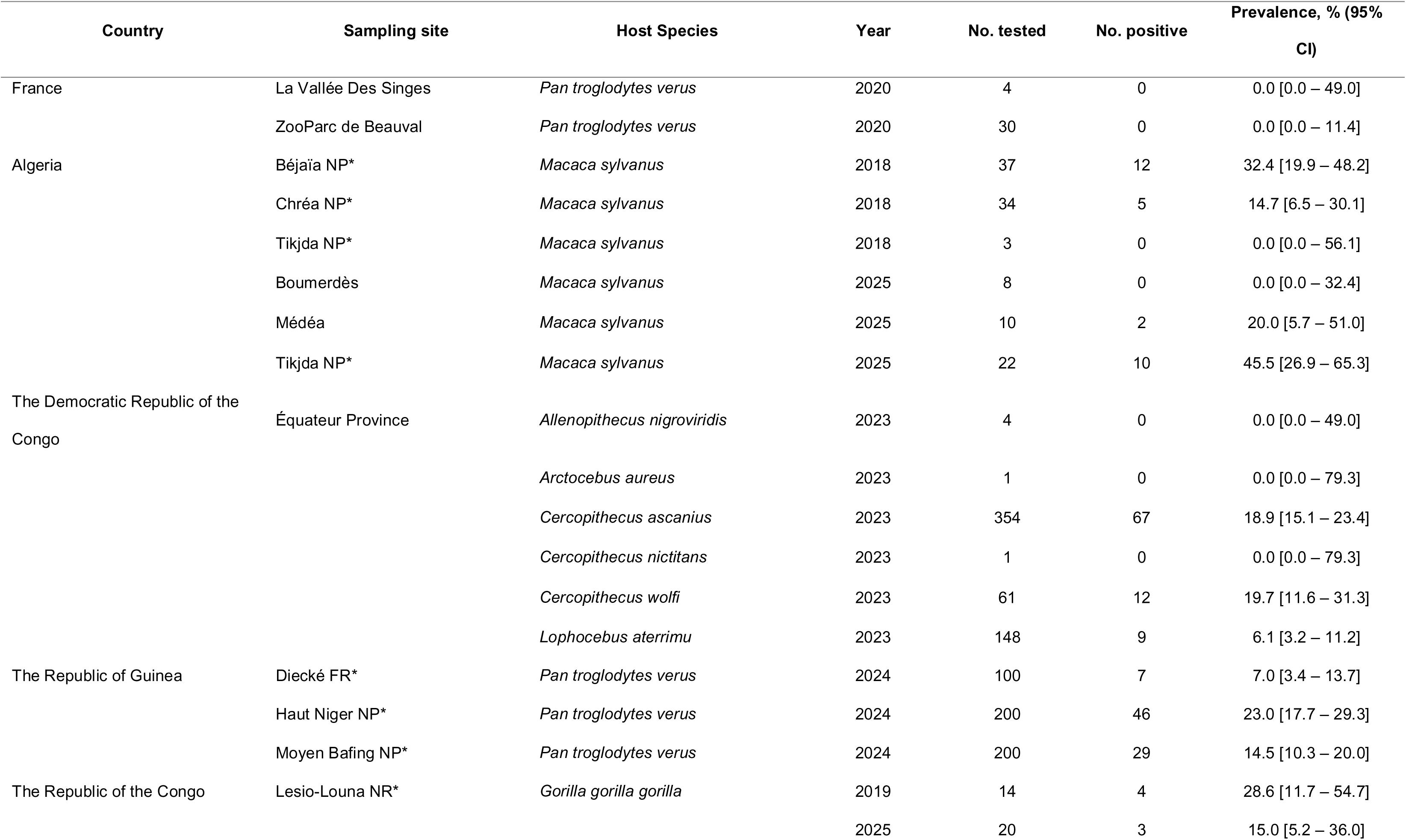
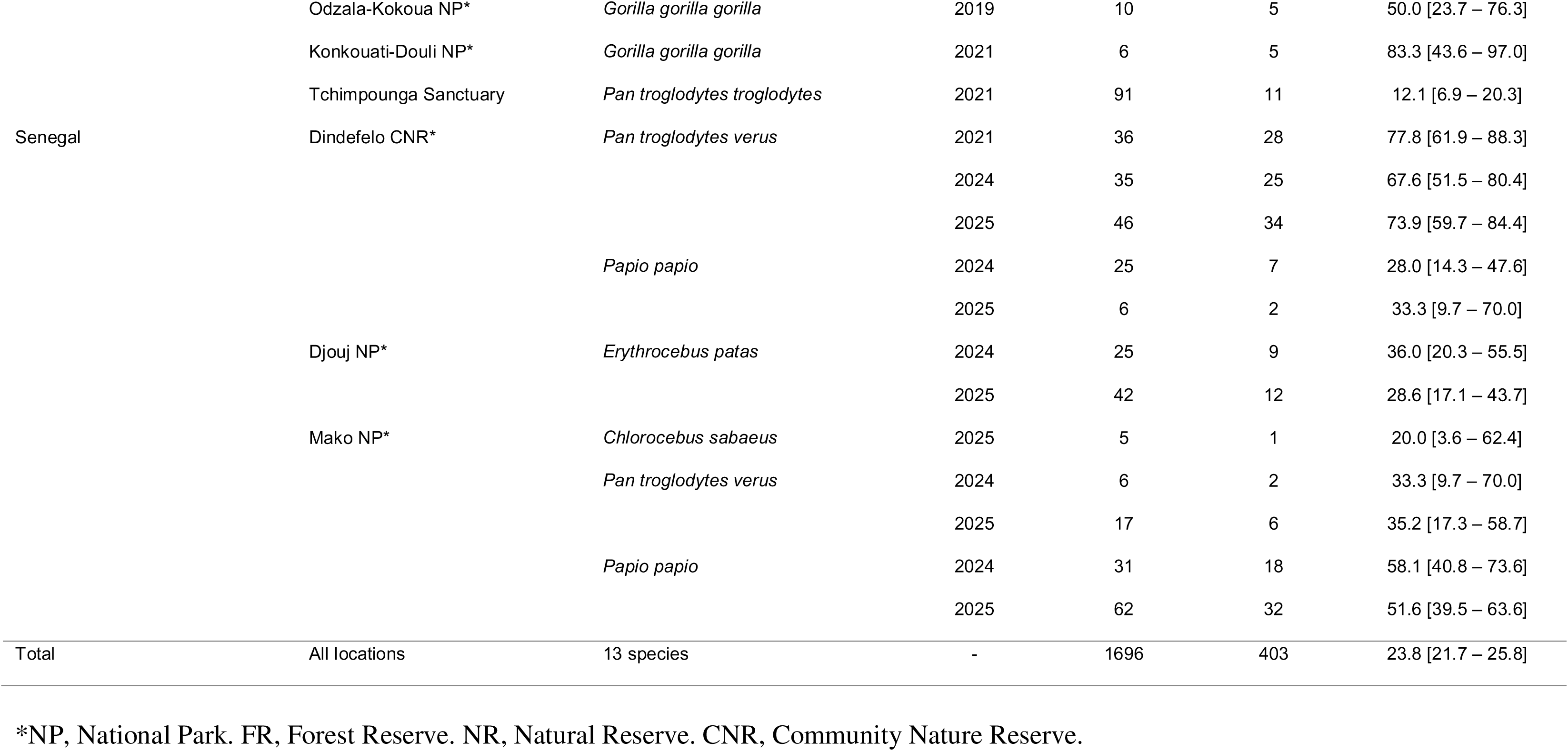
Seroprevalence of anti-*Treponema pallidum* antibodies in African nonhuman primate, 2018–2025.

### Molecular detection of T. pallidum

All 1,678 NHP fecal samples were screened for *T. pallidum* DNA using a highly sensitive real-time PCR targeting the *polA* gene. No treponemal DNA amplification was detected in any of the analyzed fecal samples. Additionally, in Senegal, despite the collection of 274 dipteran specimens belonging to *Zaprionus ghesquierei* (FJ948774), *Sphaeroceridae* sp. (KY834619) and *Drosophila yakuba* (NC001322), no samples tested positive for treponemal DNA.

### Serological detection of anti-T. pallidum antibodies

In contrast to the molecular results, widespread seropositivity was observed across all African study sites (Table 1). Detection was based on a specific chemiluminescent signal at 70 kDa, corresponding to the TpN15-TpN17-TpN47 fusion protein (Figure 2).

**Figure 2.**
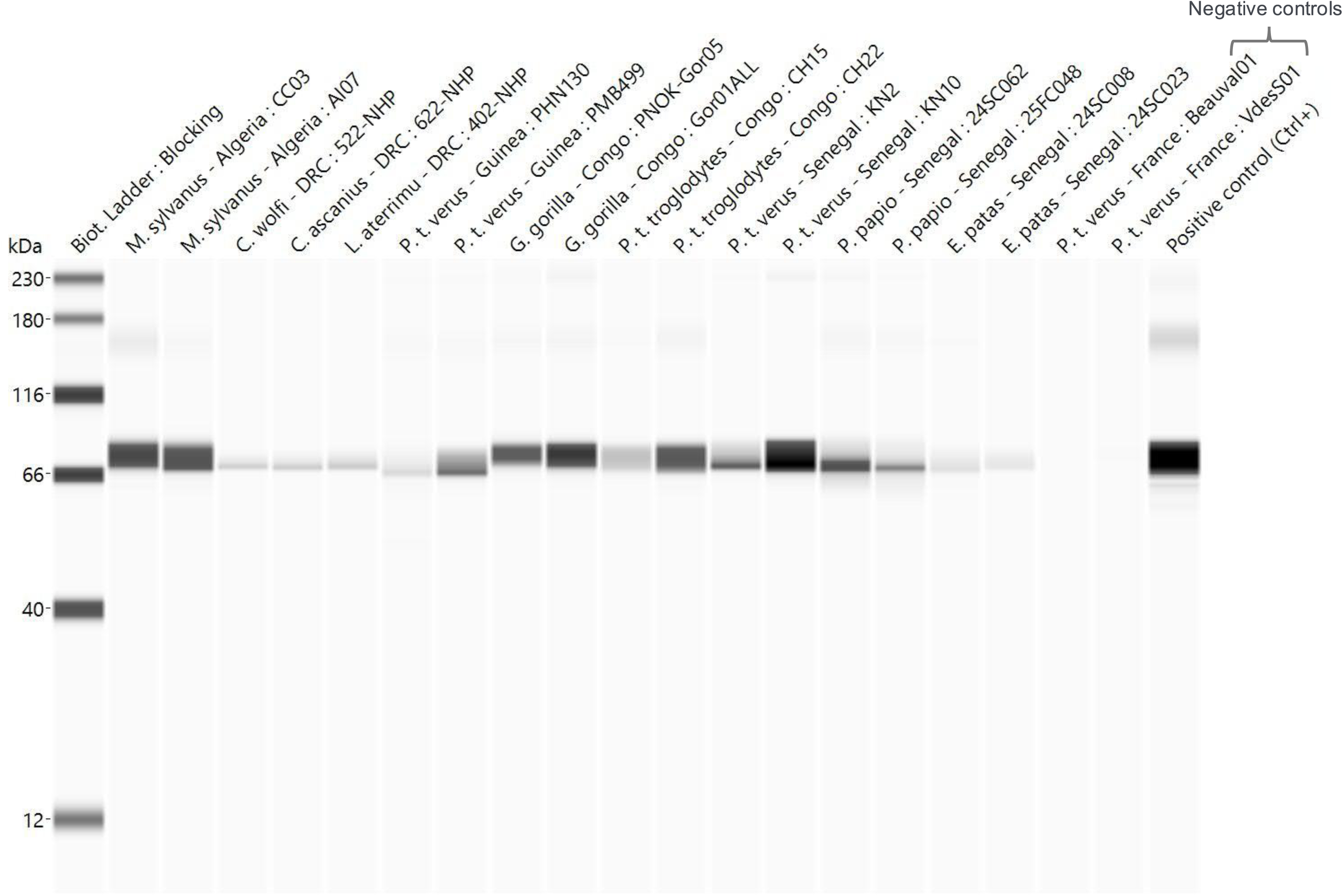
Representative automated Western blot profiles demonstrating anti-*Treponema pallidum* antibodies in African nonhuman primates fecal samples. Western blot analyses were performed using the Jess™ Simple Western platform and a recombinant 70-kDa TpN15–TpN17–TpN47 fusion protein antigen. Positive samples from *Macaca sylvanus*, *Cercopithecus wolfi*, *Cercopithecus ascanius*, *Lophocebus aterrimus*, *Pan troglodytes verus*, *Gorilla gorilla*, *Pan troglodytes troglodytes*, *Papio papio*, and *Erythrocebus patas* are shown. Seroreactivity was defined by the detection of a specific chemiluminescent signal at approximately 70-kDa corresponding to the recombinant treponemal antigen. Two nonreactive fecal samples from *P. t. verus* collected in France were included as negative controls. The lane on the far right corresponds to a human serum positive control. Molecular weight standards are indicated on the left.

The overall seroprevalence in Algeria for *M. sylvanus* reached 25.4% (29/114), with marked local variations ranging up to 32.4% (12/37) in Béjaïa (2018) and 45.5% (10/22) in Tikjda (2025). In the DRC, 18.9% (67/354) of *C. ascanius*, 19.7% (12/61) of *C. wolfi*, and 6.1% (9/148) of *L. aterrimus* samples were seropositive. *Allenopithecus nigroviridis* (0/4), *Arctocebus aureus* (0/1), and *C. nictitans* (0/1) were all seronegative. In the Republic of Guinea, an overall seroprevalence of 16.4% (82/500) was recorded among *P. t. verus* in 2024, with site-specific rates ranging from 7.0% (7/100) in Diecké Forest Reserve to 23.0% (46/200) in Haut Niger National Park. In the Republic of the Congo, 34.0% (17/50) of *G. g. gorilla* samples were seropositive, as well as 12.1% (11/91) of *P. t. troglodytes*. In Senegal, *T. pallidum* antibodies were detected across all study sites. At Dindefelo, seroprevalence remained high and stable in *P. t. verus* at 73.1% (87/119) between 2021 and 2025. In this study site, *P. papio* showed a rate of 29.0% (9/31). At Djoudj, *E. patas* exhibited an overall seroprevalence of 31.3% (21/67). At Mako, *P. t. verus* and *P. papio* showed overall positivity rates of 34.8% (8/23) and 53.8% (50/93), respectively. Finally, *C. sabaeus* at Mako showed a 20.0% (1/5) seroprevalence in 2025.

### Ecological drivers and statistical correlations of treponemal circulation

To evaluate the impact of environmental and host-related factors on pathogen circulation, sampling sites were categorized into three distinct biomes: arid/savanna (Senegal), tropical rainforest (Republic of the Congo, Republic of Guinea, and DRC), and Mediterranean (Algeria). A Chi-square test demonstrated a significant variation in seroprevalence across these biomes (χ^2^ = 84.6, *df* = 2, *p* < 0.001). Specifically, the arid/savanna biome exhibited the highest overall exposure rate (51.8%), which was significantly greater than those observed in the Mediterranean (34.4%; p < 0.05) and tropical rainforest (15.2%; *p* < 0.0001) biomes.

Furthermore, a strong correlation was identified between host taxa and seropositivity. Great apes (*P. troglodytes* and *G. g. gorilla*) were found to be 1.78 times more likely to be seropositive than cercopithecids, with respective prevalence rates of 32.4% and 21.1% ([OR] = 1.78, 95% CI [1.42–2.23], *p* < 0.0001).

Finally, longitudinal monitoring of chimpanzees at the Dindefelo site in Senegal revealed no significant fluctuations in seroprevalence between 2021 and 2025 (*p* = 0.54), suggesting a stable endemic equilibrium of *T. pallidum* within this wild primate population.

## DISCUSSION

Our study demonstrates that *T. pallidum* infection is widespread among wild NHP populations across diverse African biomes. This circulation is observed both in countries historically endemic for yaws, such as Senegal and Guinea, in regions where the disease remains endemic today, including the DRC and the Republic of the Congo, and in areas with previously undocumented epidemiological status, such as Algeria. By combining non-invasive fecal serology with a stringent cut-off methodology on the Jess^TM^ Simple Western platform, we achieved high diagnostic specificity, thereby minimizing non-specific reactivity inherent to complex fecal matrices.

We report consistently high seroprevalence of anti-*T. pallidum* antibodies across four NHP species in Senegal, confirming the region as a major endemic hotspot. Notably, infection rates in *P. t. verus* at Dindefelo (67.5% to 77.7% between 2021 and 2025) rank among the highest ever recorded in wild chimpanzees, indicating a stable and intense transmission cycle (4,10). Similarly, seropositivity in *E. patas* at Djoudj (28.6% to 36.0%) exceeds historical baseline levels reported for monkeys in northern Senegal (21). A comparable pattern is observed in *P. papio*, with seroprevalence ranging from 28.0% to 58.1%, closely mirroring data collected over six decades ago and aligning with recent field observations (3,21,22). This susceptibility is not restricted to Senegal. Across Africa, the genus *Papio* exhibits a transcontinental consistency in infection rates, suggesting that its cohesive social structure and large group sizes are primary drivers of sustained transmission (4–6,9,23).

In Guinea, where the first and only simian strain of *T. pallidum* was isolated in 1966, our results indicate widespread but overall lower transmission intensity compared to Senegal (24,25). The large sample size provides a robust representation of pathogen distribution across ecological settings. Higher prevalence in the dry wooded savannas of Haut Niger (23.0%) and Moyen Bafing (14.5%) contrasts with lower rates in the moist dense forest of Diecké (7.0%), suggesting that environmental conditions and host density differentially shape transmission dynamics.

In Central Africa, our data reveal substantial circulation of *T. pallidum* within dense rainforest ecosystems. In the DRC, seroprevalence among cercopithecids in the Équateur Province (up to 19.7%) corroborates historical observations by Fribourg-Blanc, who documented sporadic infections in chimpanzees as early as the mid-20th century (21). In the Republic of the Congo, exceptionally high infection rates were observed in gorillas (28.57–83.3%). Although based on limited sample sizes, these findings are consistent with earlier reports and recent observations from East Africa, where severe yaws-like lesions have been documented in wild *Gorilla* (26). Together, these results suggest that the forests of Congo Basin may provide optimal conditions for pathogen maintenance and transmission.

To date, no large-scale screening for treponemal infections had been conducted in North African NHPs. Our study provides the first evidence of anti-*T. pallidum* antibodies in *M. sylvanus*, with seroprevalence ranging from 30.30% to 43.33%. This macaque species has previously been screened in Southeast Asia and Gibraltar, where all tested individuals were seronegative (27). Our finding significantly expands the known geographic range of the pathogen and challenges the traditional view of treponematoses as strictly tropical diseases in

NHPs. Several hypotheses may explain the persistence of infection in this atypical biome. On the one hand, *T. pallidum* subspecies are serologically indistinguishable, and these macaques may therefore harbor a strain closely related to TEN (bejel), which was historically present in North Africa and is adapted to arid and temperate climates (2). On the other hand, these macaques could be infected with strains belonging to an ancestral lineage that colonized the Mediterranean basin during the dispersal of primates out of tropical regions. The evolutionary history of *M. sylvanus*, the only extant African macaque species and a remnant of a lineage once widely distributed across Europe, suggests a long-term host–pathogen association (28). In this context, *T. pallidum* may have persisted in North Africa following the contraction of the macaques’ historical range. These elements raise the hypothesis that North African strains could represent a distinct evolutionary lineage; however, this remains to be confirmed by genomic characterization of not-yet-isolated circulating strains. Behavioral factors, including social grooming and huddling, critical for thermoregulation in these temperate climates, may further facilitate direct intraspecies transmission (29).

Taken together, our findings support the existence of a complex continental-scale ecology of *T. pallidum* in NHPs. A key observation is the “humidity paradox”: while classical models of human yaws emphasize humid environments, our data reveal sustained transmission in arid and semi-arid ecosystems, such as the Sudanian-Sahelian savannas of Senegal and the wooded savannahs of Haut Niger. In contrast, some moist forests exhibit lower prevalence rates, such as Diecké, suggesting that humidity alone is not a determining factor. Furthermore, interspecies comparisons highlight the ability of *T. pallidum* to exploit a wide range of ecological niches. The extremely high prevalence we found in baboons contrasts with more moderate rates in the other monkey species. This disparity likely stems from differing social organizations: while baboons form large and cohesive groups and are primarily terrestrial, many cercopithecids occupy narrower arboreal niches and have smaller group sizes. Their life in the canopy, combined with smaller group sizes, may limit the intense physical interactions –such as extensive grooming or aggressive competition– that facilitate treponemal spread in ground-dwelling species with larger social groups (30,31). Thus, transmission dynamics appear shaped by a synergy of host density, sociality, and vertical stratification.

The spatial heterogeneity of clinical manifestations suggests the existence of distinct transmission routes and tissue tropisms. While clinical presentations vary across the continent, both syphilis-like genital lesions and yaws-like cutaneous manifestations have been reported across West Africa (e.g. Senegal) and East Africa (e.g. Tanzania) (3–5,8). These patterns point toward possibly multiple transmission pathways, including both sexual and non-sexual direct contact and sexual, as well as potential mechanical vectors such as flies (14,15).

However, the absence of *T. pallidum* DNA in flies collected near chimpanzee nesting sites in Senegal suggests that vector-mediated transmission, if present, may occur only under specific ecological conditions, although no necrophagous flies were recovered in this study. An additional unresolved question concerns interspecies transmission, which may result from predator-prey interactions involving infected primates, mechanical transmission by flies, or a combination of these mechanisms. A striking feature is the discrepancy between high seroprevalence and the relative scarcity of visible lesions, suggesting that latent or subclinical infections likely predominate. Nevertheless, the presence of sequelae in some individuals, including fatal outcomes in some cases, indicates that the infection can have a significant pathological impact, while the persistence of antibodies within populations reflects ongoing infection pressure.

The existence of a persistent sylvatic reservoir poses a major challenge to the World Health Organization’s goal of eradicating human yaws (13). Although zoonotic transmission has not been formally demonstrated, the occurrence of infection in primates living in close proximity to human settlements and intensive bushmeat consumption highlight a potential risk. Furthermore, as many of these host species are classified as endangered or critically endangered, the impact of treponemal infections raises important conservation concerns (32,33). Recent advances in the *in vitro* cultivation of *T. pallidum* now offer a critical opportunity to isolate novel strains, which will be essential for improving our understanding of treponemal diversity, evolution, and the mechanisms underlying differences in virulence and transmission (34). Addressing these challenges will require close collaboration among infectious disease specialists, veterinarians, and conservation biologists to assess epizootic dynamics and develop appropriate surveillance and intervention strategies.

### Conclusions

This study provides the first large-scale assessment of treponemal circulation in wild non-human primates across multiple African ecosystems. By combining non-invasive fecal serology with molecular screening, we demonstrate widespread exposure to *T. pallidum* among diverse primate taxa, spanning arid savannas, tropical rainforests, and Mediterranean environments. The absence of detectable bacterial DNA in fecal and dipteran samples, despite frequent seropositivity, suggests that infections are predominantly latent or subclinical and highlights the need for complementary approaches to investigate pathogen persistence and transmission. Our findings reveal that treponemal circulation in African primates is shaped by a complex interplay between host ecology, social behavior, and environmental context rather than by climate alone. The high and stable antibody prevalence observed in several populations, together with the identification of serological evidence in previously undocumented hosts and regions, expands the known ecological range of *T. pallidum* in wildlife. In particular, the detection of antibodies in North African Barbary macaques challenges the perception that treponematoses in non-human primates are restricted to tropical ecosystems and raises new questions regarding the evolutionary history and diversity of circulating strains. The persistence of treponemal exposure in wild primate populations has important implications for both wildlife conservation and One Health surveillance. Future studies integrating genomic characterization, clinical monitoring, and ecological investigations will be essential to identify circulating lineages, clarify transmission pathways, and assess the potential role of non-human primates as long-term reservoirs of treponemal bacteria.

## Declarations

### Ethics approval

All ethical requirements applicable to this study were fully respected. Sample collection was non-invasive and performed in accordance with institutional animal care guidelines. Fecal samples from two French zoological parks, ZooParc de Beauval and La Vallée des Singes, were collected by trained animal caretakers at the respective facilities. In Algeria, research activities and sample collections were conducted in collaboration with local authorities under authorization from the Ministry of Higher Education and Scientific Research (Directorate of Veterinary Services; N° 20/HASAQ/26; 28/06/2018). In the Democratic Republic of the Congo (DRC), sample collection were authorized by the Center for Research in Ecology and Forestry (CREF; N° 0051/MIN.R.S.I.T./CREF/MAB/DG/01/MMIK/2023; 17/11/2023). In the Republic of Guinea, research authorization was granted by the National Ethics Committee for Health Research (N°140/CNERS/24; 27/08/2024). Additionally, an official mission order was obtained (N° 000032/OGPNRF/MEDD/2024; 07/03/2024) alongside a CITES authorization (N° 005486; 11/03/2024) issued by the Ministry of Environment and Sustainable Development. In the Republic of the Congo, research authorization was granted by the Ministry of Higher Education, Scientific Research and Technological Innovation (N°003-MERSIT/DGRST/DMAST, 23/08/2023; N°05-MERSIT/DGRST/DMAST, 28/03/2024). Furthermore, access permits to national parks and protected reserves were obtained from the Congolese Agency for Wildlife and Protected Areas (ACFAP; N° 0134/ACFAP-DTS; 10/07/2018 and N° 00087/ACFAP/DG/DTS/-SRM; 19/02/2024). A CITES authorization was also provided by the General Directorate of Forest Economy, Wildlife and Protected Areas Division (N° 007; 25/08/2021). In Senegal, research authorizations were obtained from the National Parks Authority (N° 001315 and N° 001316, 22/09/2023 for the Niokolo-koba et Djoudj park during 2 years from 2023 to 2025 respectively), and the National Water, Forestry, Wildlife, and Soil Conservation Authority (N° 0003924/DEFCCS/DGF, 04/12/2020, N° 0004077/DEFCCS/DGF, 04/10/2023 and N° 0002428/DEFCCS/DGF, 28/06/2024) of Ministry of Environment and Ecological Transition. In addition, permits granting access to national parks and protected areas were obtained where required. Authorizations for the importation of animal-origin samples from non-European Union countries were also obtained for all collections.

### Consent for publication

Not applicable.

### Availability of data and materials

The datasets used and/or analysed during the current study are available from the corresponding author on reasonable request.

### Authors’ contributions

Conceptualization: CL, FF and OM. Data curation: CL, AK, CK, IA and OM. Formal analysis: CL. Investigation: CL, AK, CK, AEE, MF, GD, AB, YL, BD, IA and OM. Methodology: CL and OM. Project administration: CL, AK, CK, FF, IA and OM. Resources: CL, AK, CK, GD, AB, LD, YL, BD, RAHA, CS, FF, IA and OM. Software: CL. Supervision: FF and OM. Validation: CL, FF and OM. Visualization: CL. Writing-Original draft: CL. Writing-review & editing: all authors reviewed and edited the manuscript.

## Acknowledgments

The authors thank the Senegalese Direction des Eaux, Forêts, Chasses et de la Conservation des Sols for authorizing the collection of chimpanzee samples. We also acknowledge the Jane Goodall Institute Spain in Senegal and its team of assistants and volunteers for their logistical help in the Dindefelo Community Nature Reserve. Additionally, the authors thank the staff of ZooParc de Beauval (France) and La Vallée des Singes (France) for their technical assistance and expert advice.

